# “Micro-offline gains” convey no benefit for motor skill learning

**DOI:** 10.1101/2024.07.11.602795

**Authors:** Anwesha Das, Alexandros Karagiorgis, Jörn Diedrichsen, Max-Philipp Stenner, Elena Azañón

## Abstract

While practising a new motor skill, resting for a few seconds can improve performance immediately after the rest. This improvement has been interpreted as rapid offline learning^1,2^ (“micro-offline gains”, MOG), supported by neural replay of the trained movement sequence during rest^3^. Here, we provide evidence that MOG reflect transient performance benefits, partially mediated by motor planning, and not replay-mediated offline learning. In five experiments, participants trained to produce a sequence of finger movements as many times as possible during fixed-duration practice periods. When participants trained during 10-second practice periods, each followed by a 10-second rest period, they produced more correct keypresses during training than participants who trained without taking breaks. However, this benefit vanished within seconds after the end of training, when both groups performed under comparable conditions, revealing similar levels of skill acquisition. This challenges the idea that MOG reflect offline learning, which, if present, should result in sustained performance benefits, compared to training without breaks. Furthermore, sequence-specific replay was not necessary for MOG, given that we observed persistent MOG when participants produced random sequences that never repeated, preventing any effect of (sequence-specific) replay on the performance. Importantly, we observed diminished MOG when participants could not pre-plan the first few movements of an upcoming practice period. We conclude that “micro-offline gains” represent short-lived performance benefits that are partially driven by motor pre-planning, rather than replay-mediated offline learning.

## Results

When humans train to repeat a sequence of finger movements as often and as accurately as possible during 10-second practice periods, which alternate with 10-second rest periods, the time needed to complete a correct sequence decreases from the end of one practice period to the beginning of the next practice period. Many recent studies have interpreted these “micro-offline gains” (MOG) as a behavioral marker of micro-consolidation during rest^1–6^, supported by neural replay of the trained sequence^3,7,8^. Here, we provide evidence that MOG result neither from sequence-specific replay, nor from offline learning. Instead, MOG represent short-lived performance benefits, partially driven by motor pre-planning.

### MOG do not reflect offline learning

If MOG represent offline learning, participants should reach higher skill levels when training with breaks, compared to training without breaks. We tested this idea in an in-lab study (Experiment 1, N=85 across two groups), and confirmed results in a larger cohort via an online study (Experiment 2, N=358 across two groups, see also **Supplemental Information (SI) Figure S2 & Table S3** for an additional experiment **S1**, and **SI Figure S3** for a conceptual replication of Bönstrup et al.’s original paradigm). As in previous MOG studies^1–3^, right-handed participants trained to produce a sequence of five finger movements as often and as accurately as possible throughout fixed-duration practice periods, using their left hand (4-1-3-2-4, where “1” represents the little finger and “4” the index finger). At several time points before and during training, we assessed skill level in separate test sessions.

We assigned participants to two groups with different training schedule (**Figure 1A**). Both groups first completed a baseline test (T1) of 20 s, followed by a first block of training (Training 1). One group (‘With Breaks’) trained via three 10-second practice periods, each preceded by a 10-second rest period. In the other group (‘No Breaks’), the baseline test transitioned seamlessly into a continuous 30-second practice period without rest. In both groups, training was immediately followed by a 3-second countdown, shown on the screen, which prepared participants for the upcoming 20-second test session (T2). The countdown prompted participants to stop all ongoing movement and prepare for the upcoming test. This ensured that both groups started the test session under comparable conditions. After the test, there was a break of 5 minutes, followed by a retention test of 20 s (T3). Participants then completed a second block of training (Training 2; 30 s of practice, with or without breaks, as per group), followed by another 20-second test session (T4), and, after another 5-minute break, a final 20-second retention test (T5). Like the first test session, all following test sessions were cued on the screen by a 3-second countdown. Each of the five 20-second tests was basically a continuous practice period during which participants performed the same ‘4-1-3-2-4’ sequence as often and as accurately as they could.

**Figure 1.**
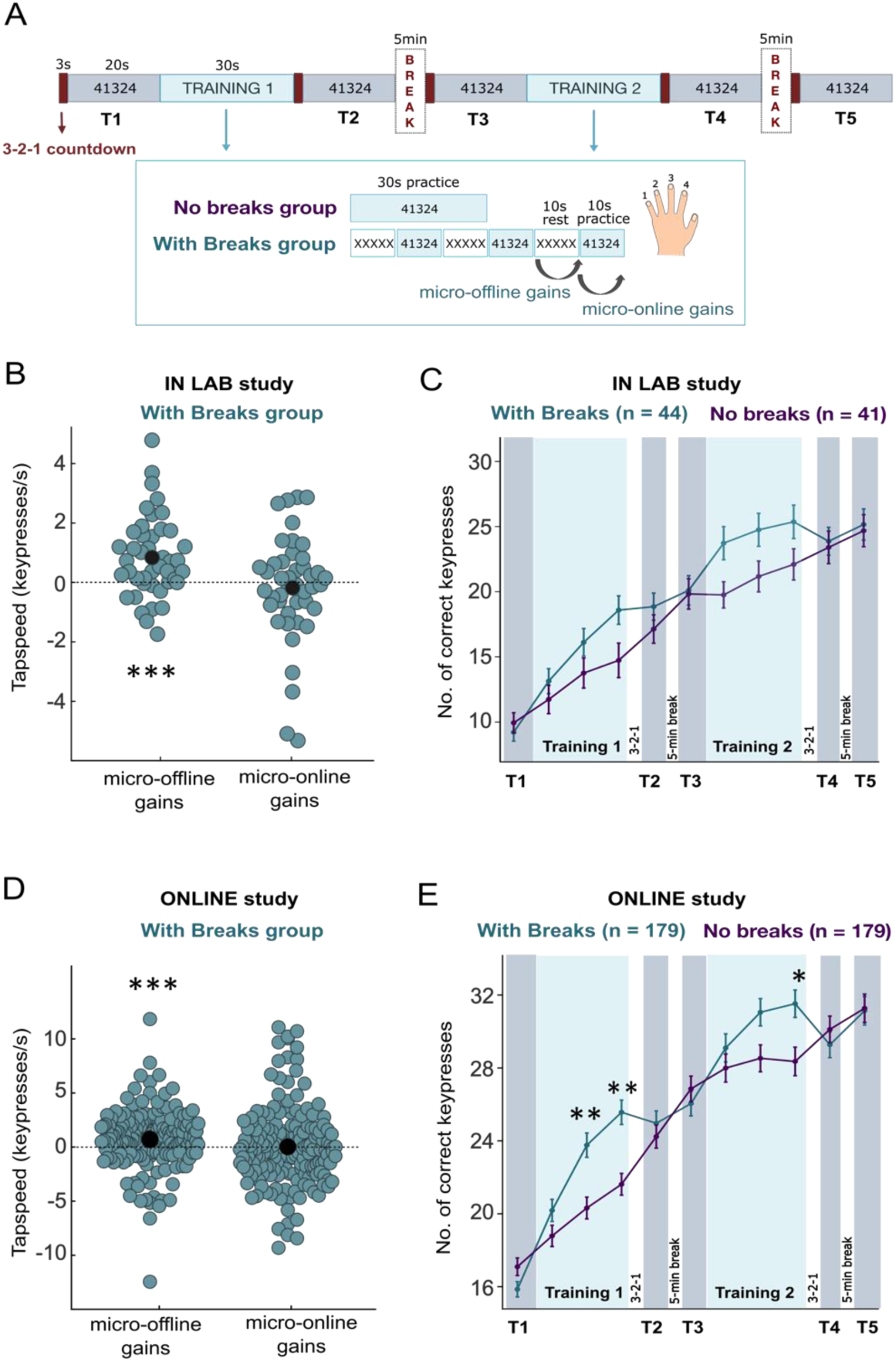
MOG do not reflect offline learning: Experiment 1 (in-lab study) and Experiment 2 (online study). (A) Experimental design. Both groups trained to produce the sequence 4-1-3-2-4 as accurately and as often as possible throughout fixed-duration practice periods, divided into two blocks of training (Training 1 and Training 2). One group of participants (‘No Breaks’) trained to produce the sequence via continuous practice periods of 30 s each, whereas the other group (‘With Breaks’) trained for the same total amount of time broken into three 10-second practice periods, interleaved with 10-second rest periods. In both groups, we assessed skill level at five time points during the experiment, via test sessions of 20 s each (T1 - T5). The task was identical for practice periods and test sessions. Panels (B) and (C) show data from the in-lab study (Experiment 1), while (D) and (E) show data from the online study (Experiment 2). Micro-online and offline were assessed for the group with breaks, as in previous studies^1,2^. ‘Micro-online gains’ are the difference in tapping speed between the last and first correct sequence within a practice. ‘Micro-offline gains’ are the difference between the last correct sequence of one practice and the first of the next. Summing these changes across 6 trials yields the total ‘online’ or ‘offline gain’ per participant. Similar results were found when computed separately for each practice (**SI Figure S1, panels F and L**). To allow for between-group comparison (panels (C) and (E)), we binned the training data into 10-second periods. For illustrative purposes, 20s test bins were also split and averaged (this also applied to the analyses of T1 and T2 when used as baseline). Error bars represent SEM. See also **SI Figure S1** for additional analyses, and **SI Figure S2** for an additional experiment.

We first confirmed that the group who trained with breaks exhibited MOG, defined identically to previous studies, i.e., faster keypresses for the first correct sequence in a 10s practice period than for the last correct sequence of the preceding practice period. We observed significant MOG both in the in-lab study (t(43)=3.922, p<.001, d=0.591, BF_10_=84.730; **Figure 1B, SI Figure S1 panel F)**, and in the online study (t(178)=3.526, p<.001, d=0.264, BF_10_=30.992; **Figure 1D**, **SI Figure S1 panel L**; one-sample t-tests against zero).

During training, performance improved more in the group with breaks than in the group without breaks. To compare training performance between groups, we binned the continuous 30-second practice periods in the group without breaks into 10-second bins. For each training block (Training 1 and 2), the preceding 20s test block (T1 and T3 respectively, averaged over two 10-second bins for each) was used as the baseline. For Training 1, a 2×4 ANOVA (two groups and four time points: baseline and three training bins) revealed a main effect of time point (in-lab study: F(2.4,199.8)=78.545, p<.001, η²_partial_=0.486; online study: F(2.9,1036.9)=189.8, p<.001, η²_partial_=0.348). In addition, there was a significant group by time point interaction (in-lab study: F(2.4,199.8)=7.604, p<.001, η²_partial_=0.084; online study: F(2.9,1036.9)=27.2, p<.001, η²_partial_=0.071). This indicates that performance improved during training in both groups, but to a different extent. In the in-lab study, planned t-tests comparing groups at each time point did not reveal any significant group differences after adjusting for multiple comparisons (all p_corr_≥.1, all BF_10_ between 1.048 and 2.049; **Figure 1C; SI Table S1**). However, in the online study, the group with breaks significantly outperformed the group without breaks at the penultimate and last time point of the first training block (both p_corr_≤ .01, BF_10_≥129, d>0.4; independent-samples t-tests; **Figure 1D; SI Table S1**). Performance during the second training block showed a similar pattern (**SI Table S1**).

Despite the performance difference during training, however, both groups ultimately reached a similar skill level: The number of correct keypresses in the two groups in all 20-second test sessions was comparable (**Figure 1C)**. A 2×5 ANOVA (two groups and five test sessions, T1-T5) indicated significant improvements across test sessions in both the in-lab study (F(2.9, 238.3)=250.075, p<.001, 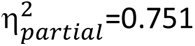) and the online study (F(3.2, 1141.4)=591.542, p<.001, 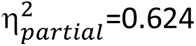). However, there was no significant group effect (in-lab study: F(1,83)=0.093, p=.761, 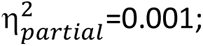 online study: F(1, 356)=0.3, p=.584, 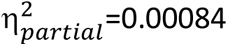). A small, but significant group x test session interaction was observed in the online study (F(3.2, 1141.4)=2.924, p=.03, 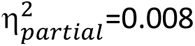), but not the in-lab study (F(2.9, 238.3)=0.776, p=.504, 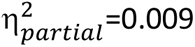). Critically, separate group comparisons for each test session revealed that there were no performance differences between groups in any test session, for both the in-lab and online studies (all p_corr_>0.6, all BF_10_≤0.35; independent-samples t-tests; detailed results in **SI Table S1**). This demonstrates that the skill level was comparable across groups, both in the test sessions completed within seconds after training (T2 and T4), and in the retention tests conducted after the 5-minute breaks (T3 and T5). Importantly, skill level was comparable across groups despite the presence of MOG in the group who trained with breaks.

In summary, taking short breaks resulted in immediate performance benefits during training (**Figure 1B-E**), as reflected in a larger number of correct keypresses produced at the end of training, and the presence of MOG. However, this benefit disappeared within seconds after training, when both groups performed under comparable conditions during test, revealing comparable skill levels in the two groups (**Figure 1C-D**). Additional analyses of several other performance metrics, including the number of correct sequences and mean tapping speed, are detailed in **SI Figure S1** and **SI Table S2** for both the in-lab and online studies. A replication of these findings in an additional experiment is presented in **SI Figure S2** and **SI Table S3**.

### MOG do not require sequence-specific replay

If MOG are supported by sequence-specific replay, as previously proposed^3^, MOG should not occur when sequences never repeat, so that any replay of previously trained sequences cannot take effect on the performance of any sequences trained later. We tested this in Experiment 3 by comparing MOG in participants who practised a single, repeating sequence ‘4-1-3-2-4’ (Repeating group, N=24 participants) with participants who produced five-element sequences that never repeated (Non-repeating group; N=19 participants; **Figure 2A**). In both groups, 10-second practice periods alternated with 10-second rest periods. In both groups we showed the first sequence of the upcoming practice period on the screen throughout the preceding rest period, such that both groups had the same level of advance information about the upcoming sequence.

**Figure 2.**
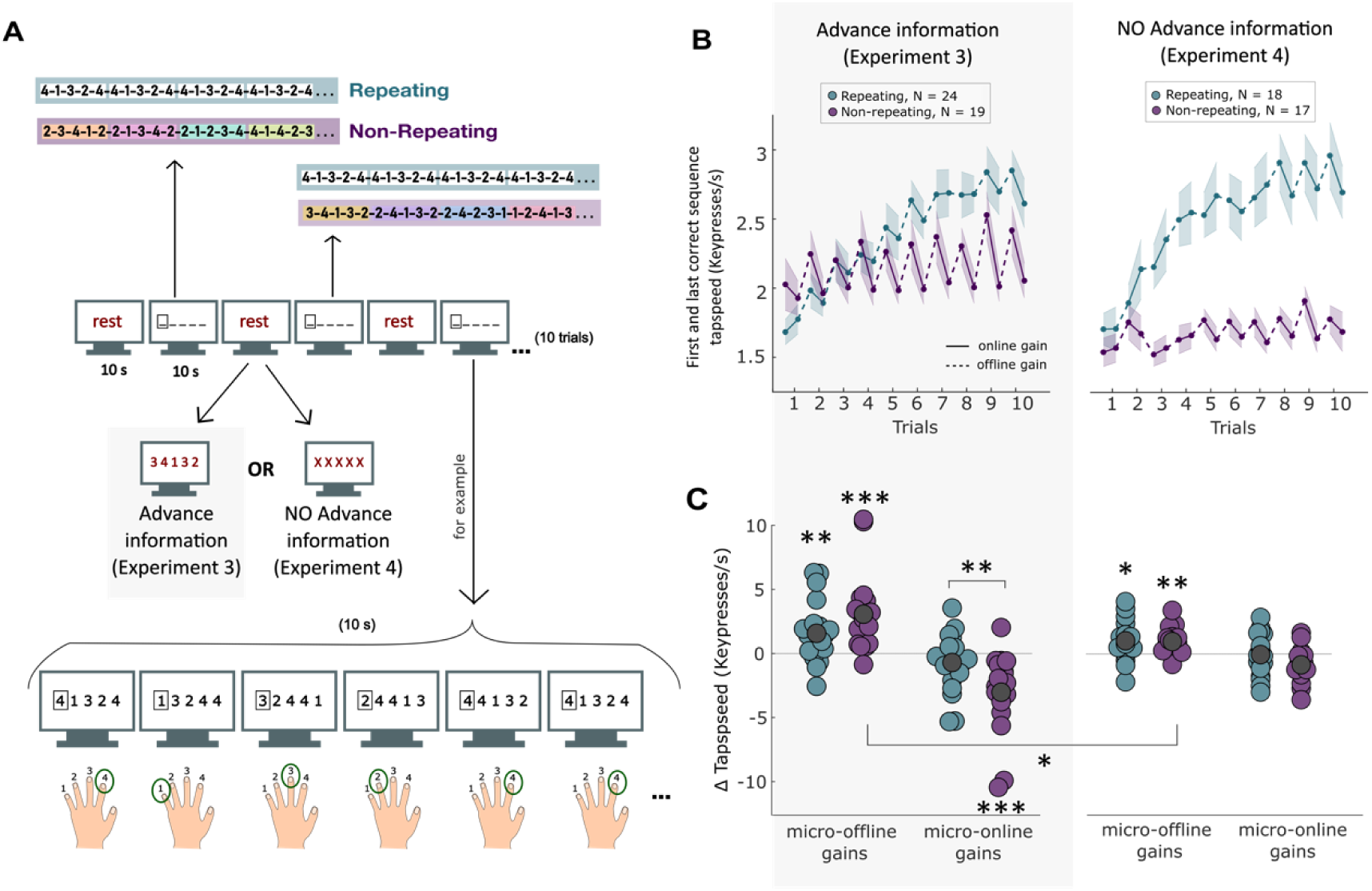
MOG do not require sequence-specific replay: Experiment 3 (advance information) and Experiment 4 (no advance information). (A) Experimental design: In 10-second practice periods, participants saw a string of five numbers, cueing a sequence of five keypresses. A rectangle surrounded the leftmost number, cueing the next movement, similar to a Serial Reaction Time Task (SRTT). Each keypress shifted the numbers leftwards, revealing a new one in the fifth position. The Repeating group saw the same 5-element sequence repeatedly (4-1-3-2-4), allowing sequence-specific replay on subsequent performance, while the non-repeating group performed non-repeating 5-element sequences, preventing reply benefits. Coloured sequence separators were for illustration; participants saw continuous strings in one colour without sequence separations. A 10-second rest preceded every practice, displaying either the next 5-element sequence (advance information, Experiment 3), or ‘XXXXX’ (no advance information, Experiment 4). (B) Tapping speed across the ten trials (10s practice periods) of each experiment. For each trial, we show the speed for the first and last correct sequence. Dotted lines indicate offline gains. (C) In both experiments, repeating and non-repeating groups showed MOG of comparable size. For the non-repeating group, MOGs were significantly larger in Experiment 3 advance information), compared to Experiment 4, where pre-planning was prevented.

As expected, performance improved more in the Repeating group than the Non-repeating group (**Figure 2B left**), as shown by a significant group x trial interaction effect on the number of correct keypresses (F(3.582,146.861)=13.745, p<.001, 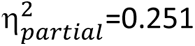). Importantly, we found significant MOG (**Figure 2C left**) in both groups (Repeating group: t(23)=3.533, p=.002, d=0.721, BF_10_=21.222, Non-repeating group: t(18)=4.515, p<.001, d=1.036, BF_10_=117.138; one-sample t-tests against zero). Furthermore, there was evidence that MOG were not larger in the Repeating group than in the Non-repeated group (t(41)=-1.902, p=.97, d=-0.584, BF_10_=0.119; one-sided, independent-samples t-test). This indicates that sequence-specific replay cannot be the cause of MOG.

Instead, the difference in learning between the Repeating and Non-repeating groups arose during ongoing practice, i.e., in ‘micro-online gains’ (**Figure 2C left**, t(41)=2.999, p=.005, d=0.921, BF_10_=9.076; independent-samples t-test). In the Non-repeating group, tapping speed declined across the 10-second practice period, following the initial post-rest performance boost that gave rise to MOG. In the Repeating group, where sequence-specific learning could offset the decline during the practice period, participants showed more even performance across the 10-second practice period. Thus, results from Experiment 3 indicate that sequence-specific learning occurred during practice, rather than in the breaks.

One possible reason for the initial performance boost is that participants in both groups could pre-plan the first few elements of the upcoming sequence before onset of the first movement. Pre-planning accelerates the first three to four keypresses of a sequence, after which execution slows down, as later keypresses need to be planned online^9,10^. Thus, pre-planning, and the resulting speed-up of the first few keypresses, could be an important driver of MOG.

### MOG partially reflect motor pre-planning

Experiment 4 tested this idea by removing any advance cueing of the upcoming sequence from the rest period (N=18 and N=17 participants in the Repeating and Non-repeating groups, respectively). While participants typically pre-plan the first elements of an upcoming movement sequence even under reaction time pressure^10^, removing the opportunity to pre-plan during the rest period should reduce MOG for the Non-Repeating group.

As in Experiment 3, performance improved more in the Repeating group than the Non-repeating group (**Figure 2B right**), as shown by a significant group x time point interaction (F(4.7,155.3)=12.058, p<.001, 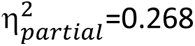). Both groups again showed significant MOG (t(17)=2.848, p=.011, d=0.671, BF_10_=4.84 for the Repeating group; t(16)=4.021, p<.001, d=0.975, BF_10_=38.26 for the Non-repeating group; one-sample t-tests against zero). As in Experiment 3, there was no difference in the magnitude of MOG between Repeating and Non-repeating groups (t(33)=0.146, p=.885, d=0.049, BF_10_=0.328; independent-samples t-test). Critically, comparing the Non-repeating groups between Experiment 3 and 4 (i.e., with vs. without advance information, **Figure 2C**), we found that MOG were reduced when the rest could not be used for pre-planning (U=248, p=.017, d=0.459, BF_10_=1.811; Mann-Whitney U test, accounting for unequal variances between the two experiment groups). This indicates that MOG are sensitive to explicit cueing of upcoming movements during rest, and, therefore, may partially reflect motor pre-planning.

Participants in the Non-repeating group in Experiment 4 could still pre-plan upcoming movements when they first saw the string of numbers that cued those movements, before the first keypress in a practice period. Such pre-planning under reaction time pressure could explain the persistent, though significantly reduced MOG in this group. Pre-planning typically delays the initiation of the first movement in a sequence^10^. Indeed, in the Non-repeating group in Experiment 4, initiation of the first movement (in practice periods that started with a fully correct sequence) was significantly delayed (884 ± 188 ms, mean ± SD), compared to the Non-repeating group in Experiment 3 (585 ± 172 ms; t(34)=4.995, p<.001, d=1.668, BF_10_=970.765; independent-samples t-test). This reaction time cost indicates that participants in the Non-repeating group in Experiment 4 may have used their reaction time to pre-plan upcoming movements, possibly accounting for the persistent MOG in this group.

We directly tested the role of pre-planning for MOG in a final, within-subject experiment that manipulated the maximum possible planning horizon (Experiment 5, N=35, **Figure 3A**). In one condition, we cued only the single next movement, so that participants could not pre-plan beyond the single next movement (‘Window size 1’, W1). In the other condition, we simultaneously cued the next five movements, so that participants could pre-plan up to five movements in advance (‘Window size 5’, W5). Participants received no advance information during rest, so that any pre-planning was restricted to the reaction time interval, i.e., the interval between the presentation of the first sequence cue and the first keypress.

**Figure 3.**
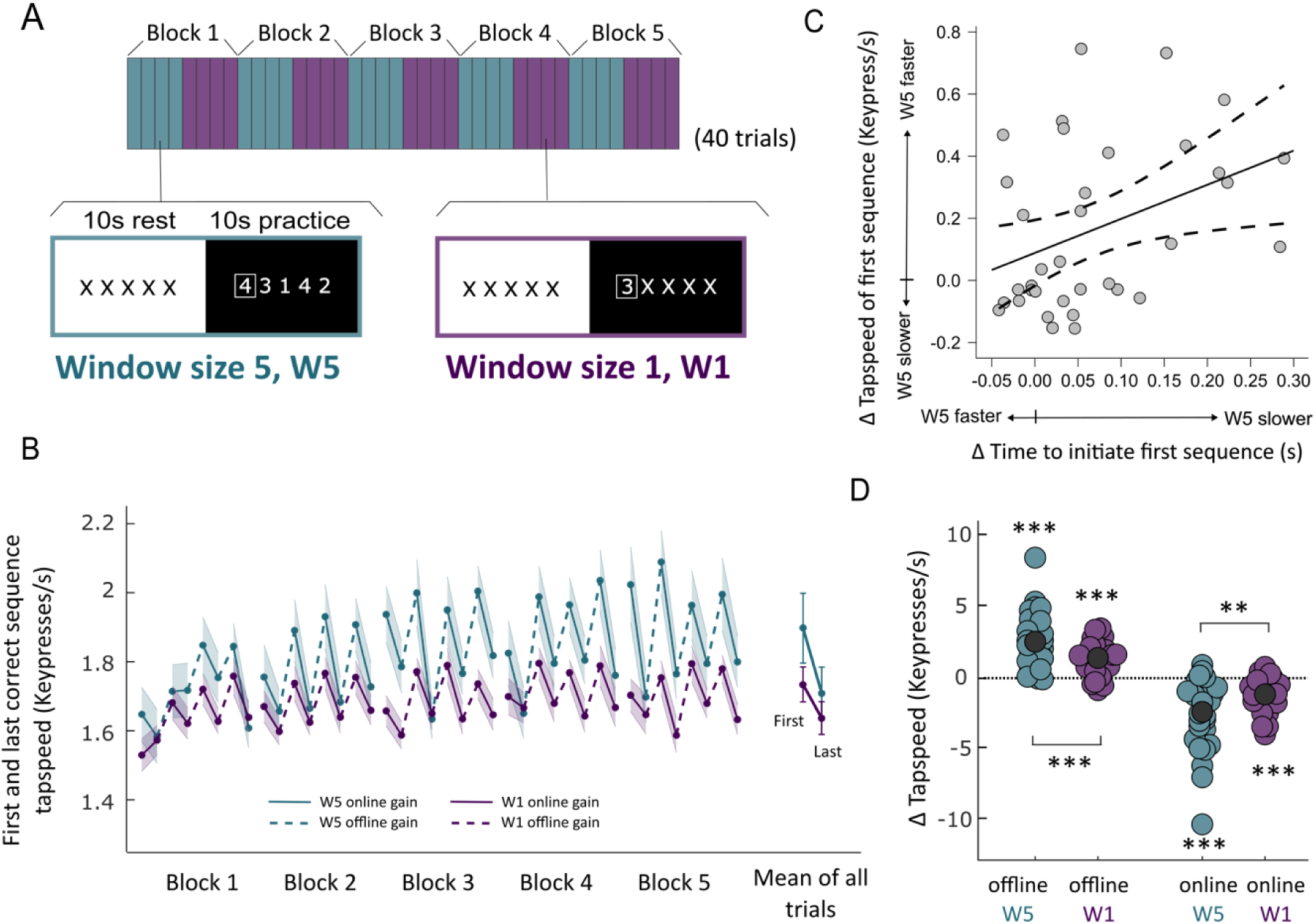
MOG partially reflect motor pre-planning: Experiment 5. (A) Experimental design. In this within-subject experiment, two conditions (Window size 5 (W5) and Window size 1 (W1)) alternated every four trials, where a trial was defined as a 10-second practice period, together with the subsequent 10-second rest period. During practice, the screen displayed a string of five numbers (W5 condition), or a string consisting of a single number followed by four X’s (W1 condition). A rectangle highlighted the leftmost number in the string (i.e., the single number in the W1 condition) as the cue for the next movement, similar to an SRTT. Across practice, participants had to perform non-repeating 5-element sequences of movements, similar to the Non-repeating group in Experiments 3 and 4. Because the string consisted of five numbers shown simultaneously in the W5 condition, participants could pre-plan up to five finger movements. This pre-planning could only occur at the start of the 10-second practice period, as each practice period was preceded by a 10-second rest period showing ‘XXXXX’. In the W1 condition, on the other hand, only the single next number was shown (and updated with every new keypress), followed by four X’s. This prevented planning of more than a single finger movement at a time. (B) Increasing the window size (from the W1 condition to the W5 condition) resulted in performance benefits that were stronger for the first correct sequence than for the last correct sequence. (C) Consistent with pre-planning, participants traded-off the speed of initiating the first movement in a practice period, and the tapping speed of the first correct sequence. Participants with slower movement initiation in the W5 condition, compared to the W1 condition, were faster in executing the first correct sequence in the W5 condition, compared to the W1 condition, and vice versa. (D) We found significant MOG in both conditions, however, MOG were significantly smaller in the W1 condition. This confirms that pre-planning contributes to MOG.

The pattern of initiation time for the first movement after a break validated our experimental manipulation. Consistent with pre-planning, initiation times for the first movement after a break were significantly slower in W5 trials (796 ± 126 ms, mean ± SD) compared to W1 trials (728 ± 122 ms; t(34)=4.343, p<.001, d=0.734, BF10=212.874), considering only those practice periods that started with a fully correct sequence). This delay allowed them to be faster in executing the first 5-element sequence. A 2×2 ANOVA, comparing tapping speed of the first vs. last correct sequence of a practice period in the W5 and W1 conditions, provided evidence that a larger window size improved performance in particular at the for the first sequence of a practice period, reflected in a significant condition x position interaction effect (F(1,34)=12.111, p=.001, 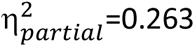). While tapping speed was faster for the W5 compared to the W1 condition for the last correct sequence (t(34)=2.415, p_corr_=.021, d=0.183, BF_10_=2.268), this effect was more pronounced for the first correct sequence (t(34)=3.661, p_corr_<.001, d=0.619, BF_10_=36.872; **Figure 3B**). Furthermore, participants who showed pronounced initiation time costs in the W5 condition (i.e., took longer to initiate the first response) also displayed a larger benefit in execution time of the first sequence (r=.385, p=.022; considering only trials in which the first five keypresses were correct; **Figure 3C**).

As a consequence, MOG were significantly larger in the W5 condition than in the W1 condition (t(34) = 3.686, p<.001, d=0.623, BF_10_=39.249; **Figure 3D**). Preventing pre-planning beyond the single next movement, however, did not completely abolish the slowing during the 10s practice period. As such, we observed significant MOG in both conditions against zero (t(34)=8.237, p<.001, d=1.392, BF_10_=8.876x10e6, W5 condition; t(34)=7.849, p<.001, d=1.327, BF_10_=3.159x10e6, W1 condition).

## Discussion

Our results challenge the current view that MOG reflect offline learning^1–6^ mediated by sequence-specific replay^3,7^. We found that training with breaks gives rise to MOG, but conveys no lasting learning benefit over training without breaks, casting doubt on the idea that MOG result from additional (offline) learning during breaks. Furthermore, MOG also occurred for non-repeating sequences, indicating that they are not tied to the replay or consolidation of a specific motor memory. Instead, our data indicate that MOG represent short-lived performance benefits that are at least partly mediated by pre-planning.

Introducing breaks between practice sessions (spaced training) can enhance long-term learning success in many tasks, compared to massed training. However, while there is extensive research on spacing effects on declarative knowledge^11^, from vocabulary learning^12^ to mathematics^13^ (for a review, see Smolen et al.^14^), the effects of spacing on motor skill acquisition^15–19^, particularly on motor sequence learning^18^, has remained largely unexplored. While spacing improved performance during training in our study, this benefit rapidly dissipated in test sessions that followed within seconds after training. These test sessions allowed for an unbiased comparison of skill levels, and revealed that people acquire similar skill levels whether, or not, they train with breaks. Our findings, therefore, do not support a spacing effect in motor sequence learning on a short time scale in the order of seconds. Instead, they highlight a crucial distinction in motor skill learning, as noted by Kantak et al.^20^, between motor performance, i.e., observable changes during training, and motor learning, i.e., the longer-term retention and stability of skills developed through training^20–22^.

Consolidation of declarative and spatial memories is known to involve hippocampus^23–26^. Buch et al.^3^ found that MOG correlated with rapid replay of the trained sequence during rest periods in magnetoencephalography (MEG) data source-localised to hippocampus (see also^7^). If neural replay of the trained sequence were driving MOG, MOG should vanish when sequences never repeat during training, because this would prevent any (sequence-specific) effect of replay on subsequent performance. Yet Experiments 3 and 4 demonstrated that MOG were of similar size even for sequences that never repeated. While our findings do not rule out neural replay during rest, they clearly show that sequence-specific replay is not necessary for MOG to occur. Consequently, interpretations of MOG as a behavioural index of replay-mediated micro-consolidation – as extensively discussed in the literature^3,5,6^ in relation with beta band activity^1^ or with hippocampal ripples^7,8^ – should be considered with caution.

Blood oxygenation level dependent (BOLD) signals in the hippocampus and precuneus during rest correlate with MOG^5^. Additionally, learning-related patterns in functional MRI persist from training into short inter-practice rests^6^. While these findings have been interpreted as indicative of a memory reactivation of the sequence trained in the past, they may alternatively reflect planning of future sequence production. Kornysheva et al.^27^ identified an MEG signature of motor sequence planning that involved medial temporal lobe, and that linked practice-related information with the preceding planning stage, broadly consistent with the abovementioned fMRI studies. They found that several finger movements in an upcoming sequence could be decoded in parallel during preparation. This parallel planning signature source-localized to the parahippocampus and cerebellum, and predicted motor sequence performance^27,28^.

Rest interspersed with training could thus provide an opportunity to plan the first elements of the upcoming sequence before re-starting training. In that regard, the disruptive impact of theta burst stimulation of the dorsolateral prefrontal cortex (DLPFC) on the degree of MOG^29^ may potentially be attributed to the DLPFC’s critical role in sequential event planning^30,31^. Building on this, in Experiment 5 we observed diminished MOG when the rest period, as well as the time before first movement initiation, could not be used for planning several upcoming finger movements, compared to conditions allowing for pre-planning. Thus, pre-planning contributes to MOG.

In the non-repeating group of all of our studies, we observed a clear slowing of performance during each of the 10-second practice periods (Figure 2B, 3B). Such slowing could arise due to a switch from fast pre-planned movements to slower online planning^9^ and / or fatigue (with motor slowing observed already within 10 s^32^ or reactive inhibition^33^. When analysed through the lens of online- and offline-gains, the slowing leads to “negative micro-online gains” for the non-repeating group.

The same slowing also induces a strong bias when using the current methods to asses MOG: When using the difference in speed of the first correct sequence performed after the rest to that of the last sequence before the rest to measure “offline learning”, we need to assume that the performance is stable throughout the practice period. If performance declines over time during the practice period, this approach will underestimate online learning and overestimate improvements imparted by the rest period.

For the repeating group, the slowing across the 10s of practice will combine with actual learning during the practice period, often resulting in no net performance change from the beginning to the end of a practice session. In our experiment, the online gains for repeating group in Experiment 3 were close to zero. Similarly, other previous studies have found no net change in performance during 10-second practice periods ^1,3,5^, while other studies have found a net slowing of performance ^7^, as in our conceptual replication of Bönstrup et al.’s paradigm (**SI Figure S3**).

Our finding of a difference between repeating and non-repeating groups in online, but not offline, gains, provides clear evidence that the learning actually occurred during the practice periods, but was masked by an additive slowing effect. Subsequently, the latent effects of learning during practice then become evident after the rest period, after which pre-planning can occur again, or after fatigue^32^ and reactive inhibition have dissipated^33,34^. Thus, sequence-specific learning actually occurred during the practice periods, which then became unmasked after the rest, providing a temporary performance boost (equivalent for Repeating and Non-repeating groups).

In conclusion, we propose that the improved performance observed after a short break are short-lived and do not depend on sequence-specific neural replay. The cause of the post-rest performance improvements are likely multifactorial, with the ability to pre-plan movements playing an important role. Other contributing factors could include the dissipation of reactive inhibition or fatigue during rest. Moreover, attentional recovery facilitated by context switching^35^, in conjunction with pre-planning, may explain why a pause of only 3 seconds is sufficient to unveil true performance levels.

## Acknowledgements

We thank Ben Ford, Kamil Kilic, and Rosanna Franke for support with programming and data collection. M.-P.S. and E.A. received funding from the German Research Foundation (SFB-1436, TPC03, project-ID 425899996), which supported the PhD positions of A.D. and A.K. J.D. was supported by the Canada First Research Excellence Fund (BrainsCAN). M.-P.S. received funding from the Volkswagen Foundation via a Freigeist Fellowship (project-ID 92977).

## Author contributions

**Conceptualization** EA, MPS, JD

**Methodology** AD, EA, MPS, JD

**Software** AD, EA, MPS

**Validation** AD, EA, MPS, AK, JD

**Formal Analysis** AD, EA, MPS, JD

**Investigation** AD, EA, MPS, AK

**Resources** EA, MPS

**Data Curation** AD, EA, MPS

**Writing – Original Draft** AD, EA, MPS

**Writing – Review & Editing** AD, EA, MPS, JD, AK

**Visualization** AD, EA, MPS, JD

**Supervision** EA, MPS, JD

**Project Administration** AD, EA, MPS

**Funding Acquisition** EA, MPS

## Resource availability

Corresponding authors: For further information and resource requests, please contact **Anwesha Das** or **Max-Philipp Stenner**.

## Methods

Across the five experiments, a total of 631 participants (270 females) took part in this study. 89 (31 females, mean age = 25.9 years, SD = 3.2 years) took part in Experiment 1 and 413 (175 females, mean age = 27.9 years, SD = 7.7 years) took part in Experiment 2, 49 (29 females, mean age = 39.9 years, SD = 15.8 years) in Experiment 3, 37 (20 females, mean age = 39 years, SD = 13.5 years) in Experiment 4 and 43 (15 females, mean age = 27.3 years, SD = 3.2 years) took part in Experiment 5. For Experiment 1, we calculated the sample size sufficient to detect a difference in the number of correct keypresses between two groups of participants (with and without breaks) that is at least medium-sized (Cohen’s d≥0.6) with at least 80% power (independent-samples t-test; the pooled standard deviation was estimated from a previous study with 62 participants (see **SI Figure S2**)). In Experiment 2, our goal was to achieve ≥80% power to observe an effect of group on number of correct keypresses of d≥0.3. Experiments 3 and 4 were crowdsourcing studies during a science communication event at the Leibniz Institute for Neurobiology (the Magdeburg Long Night of Science), during which we collected data from as many volunteers during the six hours of the event as possible (total of 86 adult datasets collected). Cohort size for Experiment 5 was determined by our goal to observe any effect of pre-planning on ‘micro-offline gains’ of size d≥0.5 with ≥80% power.

Participants of all experiments, except the crowdsourced experiments (Experiments 2, 3 and 4), were recruited based on the following exclusion criteria. They had to be right-handed, between 18 and 40 years of age, could not be professional typists or skilled musicians (i.e., recruited participants did not play any musical instrument requiring skilled finger movements for consecutive 4 years at any point in their life), did not have any prior or existing neurological or psychiatric conditions, and were naïve to the task. Participants for Experiment 2 were recruited via the following criteria. They had to be right-handed, between 18-40 years of age, and without prior neurological or psychiatric condition. There were no criteria applied for participation in Experiments 3 and 4, but we performed analysis only on the data of participants who were right-handed and 18 years of age and above. Participants for the in-lab studies were recruited via local participant databases (Sona systems, https://magdeburg.sona-systems.com/) whose members are mostly students and staff of Otto-von-Guericke University in Magdeburg, whereas participants for the online study (Experiment 2) were recruited via the Prolific database (www.prolific.com). Participants for Experiments 3 and 4 were volunteers during a science communication event, as described above.

Handedness was determined by the Edinburgh Handedness Inventory assessment (Oldfield et al^36^). The study was approved by the ethics committee of the Otto-von-Guericke University Magdeburg, Germany, and conducted in accordance with the Declaration of Helsinki. All participants, except for visitors of the Long Night of Science taking part in Experiments 3 & 4, received financial reimbursement for their time of participation, and were additionally rewarded with bonus money based on their performance in the task.

### Apparatus

All experiments except the online crowdsourced study were conducted in a behavioral laboratory (a room with four computers on desks, and chairs arranged in the form of cubicles, for participants to sit and perform the task in privacy without distraction) using LCD displays, PC keyboards and headphones. Stimuli for Experiments 1 and 3-5 were presented using MATLAB (R2021b, The MathWorks, Inc., Natick, Massachusetts, United States) and Psychtoolbox^37^, whereas Experiment 2 (the online study) was programmed in PscyhoPy^38^ and run on Prolific (www.prolific.com). In the laboratory, we tested either single participants, or up to four participants at a time simultaneously.

### General task design

Participants were asked to sit in front of an LCD monitor (60 Hz) and place the little finger, ring finger, middle finger, and index finger of their left hand on four keys of the keyboard. Throughout all experiments except Experiment 2, we used the keys F, T, Z and J (on a German-layout QWERTZ keyboard). In the (online) Experiment 2, participants were instructed to use the numbered keys 1, 2, 3, 4 located on top of the keyboard. These keys were uniquely associated with the numbers 1 to 4 displayed on the monitor, so that the number 1 (or key F) corresponded to the little finger, 2 (or key T) corresponded to the ring finger, 3 (or key Z) corresponded to the middle finger, 4 (or key J) corresponded to the index finger. In Experiments 1 and 2, the screen displayed a static string of five numbers (4-1-3-2-4), for a certain duration depending on the experiment design. The task was to produce that sequence using the corresponding fingers as fast and accurately and as many times as possible throughout the practice period, i.e., as long as the sequence stayed on the screen. The string of numbers was displayed in white on a black background, surrounded by the outline of a rectangle, whose color changed with every keypress, from white to grey, or back from grey to white, as feedback that the keypresses were being registered. In Experiments 3, 4 and 5, only the first number of the sequence was surrounded by a white square, and the display of the five numbers changed with every keypress, replacing the first number with the second, the second with the third, and so on, and adding a new number as the fifth. This was done to accommodate non-repeating finger sequences, as described in detail in the corresponding section below. In the different experiments, the durations of the practice periods changed depending on the experimental manipulation, but the task instruction for practice periods remained the same throughout.

The practice periods were interleaved with rest periods of 10-seconds in some experiments. In Experiments 1, 2, 4 and 5, the screen displayed a static string consisting of five times the letter X (i.e,, ‘XXXXX’) during rest periods, replacing the numbers. Participants were asked to fixate on the X’s while resting, and not move their fingers or any other body part. In the rest periods of Experiment 3, participants were shown the first sequence of the upcoming practice period, in red font (see details below).

### Experiments 1 & 2 (with versus without breaks, in-lab and online study)

In order to test if offline periods during training resulted in additional learning, i.e., overall more skill acquisition, Experiments 1 and 2 followed a between-subject design with two groups of participants: one group trained with interspersed breaks (‘With Breaks’) whereas another group trained continuously for the same duration without interspersed breaks (‘No Breaks’). We ran Experiment 1 with a total of 85 participants tested in-lab, and Experiment 2 with a total of 358 participants who participated online. During training, participants in the ‘With Breaks’ group had to learn the sequence ‘4-1-3-2-4’ via 10-second practice periods, interspersed with 10-second rest. Participants in the other group (‘No Breaks’) had to learn the same sequence via continuous practice without breaks. The total practice duration was matched between groups. Following training, both groups took a break of 5 minutes, which allowed for washout of fatigue. Test sessions of 20 s each were introduced before the beginning of training, at end of training, and at the end of washout (**Figure 1A**). The task during these test sessions was identical to the practice periods. In order to explore potential group differences across longer training, a second block of training, washout, and tests was added. Experiment 2 (online) had the identical task design, with the exception that two attention checks were introduced during each of the 5-minute washout periods, in order to ensure that the participants did not move away from the keyboard. Each attention check required participants to press a key within the number of seconds shown on the screen. Participants in both experiments were motivated to perform to the best of their capacity by rewarding them bonus money based on the total number of correct sequences they performed throughout the experiment.

### Experiments 3 & 4 (Repeating versus non-repeating sequences, with and without advance information)

Experiments 3 and 4 tested whether the performance improvement across rest requires replay. To this end, one group (‘Non-repeating’) each in Experiments 3 and 4 performed sequences of finger movements which never repeated, neither within a given practice period, nor across the entire experiment. If MOG are due to replay, they should vanish when sequences never repeat. This is because replay of any sequences that were previously encountered can only improve future performance of that same sequence, i.e., benefits mediated by this type of specific replay effects would require that sequences repeat across practice periods. To accommodate never-repeating sequences, we modified the practice periods to closely mimic a DSP^9^ (discrete sequence production) task. Every practice period was 10 s in duration. During that time, participants always saw five numbers at a time on the screen (in white on a black background). The first number was surrounded by a square frame and cued the next required movement. Participants were instructed to press the key corresponding to that number as accurately and fast as possible. As soon as they pressed, the sequence of numbers moved to the left by one position, such that the number within the square disappeared, and the second number from the left moved into its position, while a new number appeared in the fifth position. Practice periods lasted 10 seconds, so that the participants paced the rate of cue presentation. The ‘Repeating’ group of participants performed the same 5-element sequence of numbers (4-1-3-2-4) repeatedly, whereas the ‘Non-repeating’ group of participants were presented with a set of 5-element sequences that never repeated. Both groups completed 10 trials of alternating practice and rest periods. To mimic the 4-1-3-2-4 structure of the repeating sequence, the non-repeating sequences were generated with some constraints: all four possible numbers (1-4) had to be present, with any one of these numbers presented twice in a sequence of 5 elements, and consecutive repetitions of a single number or of a pair of numbers within the sequence were not allowed. These unique 5-element sequences were organised into a long string such that the last number of a sequence was the same as the first number of the next sequence (e.g., ‘<u>2-4-2-3-1</u>-<u>1-2-4-1-3</u>…’), in order to match the Repeating group (‘<u>4-1-3-2-4</u>-<u>4-1-3-2-4</u>…’). Participants of neither group were explicitly aware of the sequence demarcations.

In Experiment 3, both groups received advance information about the first sequence of an upcoming practice period, keeping it similar to the original studies^1,3^, in which participants knew throughout rest periods which sequence they would have to perform next. Therefore, the rest periods displayed the first five cues of the upcoming practice period, in the form of a 5-element sequence displayed in red. For the ‘Repeating’ group, this sequence was always 4-1-3-2-4, and for the ‘Non-repeating’ group, it was the first sequence of the upcoming practice period.

In Experiment 4, on the other hand, the rest period preceding every practice period displayed ‘XXXXX’ in red at the centre of the black screen. This restricted participants from pre-planning the first sequence during rest.

### Experiment 5 (pre-planning)

In this within-subject study, we tested if preventing planning of more than the next finger movement diminishes MOG. All the sequences in all trials were non-repeating. There were 40 trials, each having a 10-second rest period and a 10-second practice period, with two interleaved conditions: ‘Window size 5’ and ‘Window size 1’, which allowed to pre-plan up to 5 upcoming movements or only the single next movement, respectively. The two conditions switched every four trials. In the ‘Window size 5’ condition, the practice period showed a sequence of five numbers in white, with the leftmost number being surrounded by a white rectangle. Thus, participants were able to know the full sequence and pre-plan up to five finger movements before initiating the first finger movement at the start of each practice period, as well as plan online. Participants were instructed to respond as correctly and as quickly as they could, by pressing the key corresponding to the number within the rectangle, using their left hand’s corresponding finger. As soon as they pressed, the sequence shifted one number to the left, as described for Experiments 3 and 4. Participants were instructed to respond to the number within the rectangle at all times. In the ‘Window size 1’ condition, the practice period showed a single number within the white rectangle, followed by four X’s to its right (for example, ‘3 X X X X’), thus removing the possibility for planning any movement beyond the single next movement. Participants were instructed to respond as correctly and immediately as possible. As soon as they pressed a key, the white rectangle showed the next number to be pressed, while the X’s stayed in the same position. All presented numbers in both conditions belonged to 5-element sequences, unknown to participants. All practice periods were timed to 10 seconds, so the number of sequences that each participant encountered depended on their speed of responses. Sequences were generated in a way that there was no single number, or pair of numbers, that appeared consecutively within the same sequence. We also ensured that all four fingers were represented in each sequence at least once, and that any one finger number appeared twice in the combination in order to obtain a 5-element sequence. Each practice period was preceded by a rest period. During rest, participants saw ‘XXXXX’ in white on a black background, and were asked to fixate on the ‘XXXXX’ without making any movement, and rest their fingers on the keys. Thus, any pre-planning in the ‘Window size 5’ condition could only occur once the practice period began and the five numbers were displayed.

### Data analysis

#### Dataset exclusion criteria

Out of the complete datasets collected for each experiment, the following number of datasets were discarded from analysis for the respective experiments (see Exclusion Criteria table below): 4 from Experiment 1, 55 from Experiment 2, 8 from Experiments 3 & 4, and 8 from Experiment 5.

In Experiment 2, all participants’ data were screened to check for completion, attention tests and time of initiation after the washout periods. Fully complete datasets which passed both attention tests for both washout periods, and showed initiation times less than 2.5 s from the start of the post-washout test, were included for analysis.

| Experiment | Reason for exclusion (No. of participants excluded) |
| --- | --- |
| Experiment 1 (breaks vs no breaks): in-lab study | Anomalous performance - Zero correct keypresses after baseline and 1st training (1)<br>Anomalous performance - Zero correct keypresses in the last test T5, although they performed well throughout (1)<br>Problems during data acquisition - incorrect keyboard, failure to follow instructions (2) |
| Experiment 2 (breaks vs no breaks): online study | Failed attention checks during 5-min breaks (5)<br>Poor task commitment - took more than 2.5 s to make first response after task started post-5min breaks (38)<br>Unable to follow task instructions – pressed wrong keys in a pattern throughout the experiment (12) |
| Experiments 3 & 4 (Repeating vs Non-repeating sequences, with & without advance information) | Due to poor commitment to the task – zero correct sequence performed in multiple trials (6 & 2) |
| Experiment 5 (pre-planning) | Due to technical issues during data acquisition leading to invalid datasets (7)<br>Due to poor performance, pressing wrong keys on the keyboard (1) |

All recorded data were stored anonymously and with the consent of participants under GDPR regulations. Information about identity and timings of the keypresses were recorded, for every practice and rest period in all the experiments. Most of the data analyses were done with the same methods as used by previous studies^1–3,5,39^.

#### Number of correct keypresses

While the task required producing as many fully correct sequences as possible during each practice period, errors likely resulted in sequences that were only partially correct (for example, four of the five required keypresses correct). Quantifying performance purely as the number of fully correct sequences would therefore disregard part of the task-appropriate performance, i.e., sequences that were only partially correct. To avoid this, we computed the number of correct keypresses per practice period, rather than the number of fully correct sequences^1^. Our approach to calculate the number of correct keypresses differed between Experiments 1 and 2 on the one hand, and Experiments 3-5 on the other hand. In Experiments 1 and 2, the same sequence was required throughout the entire experiment, and we did not cue single keypresses, one at a time. As a result, when they made an error, participants were free to complete the current sequence despite the error, or restart. To accommodate this, we used a sliding window approach. We first calculated keypresses that were part of fully correct sequences. For calculating the number of keypresses which were not part of a fully correct sequence, the positions of all keypresses in fully correct sequences were replaced by NaN (as a placeholder), and we then moved a 5-element sliding window to match keypresses in the current window with the required sequence position-wise, according to the following criterion. If at least three keypresses in the current 5-element window matched the required sequence by position, these keypresses were counted as correct keypresses, and they were replaced by NaN before the sliding window moved on. The reason why we set the threshold to at least three correct keypresses was to only count correct attempts to perform a fully correct sequence, and disregard instances when participants repeated, e.g., the first one or two elements of the sequence. In the case of Experiments 3, 4 and 5, in which keypresses were cued individually, one at a time, all correct responses to the currently valid cue were counted as correct keypresses.

In Experiments 1 and 2, the continuous training durations of the group with no breaks were binned into three 10-second bins, for comparison with the group with breaks. The number of correct key presses was calculated in the same manner as for the 20-second bins, as explained above. Thus, the sliding window used for detecting correct keypresses matched keys to the sequence based on their position. As a result, it could miss detecting the first keypresses of a bin, if the bin did not start with a correct ‘4-1-3-..’ initiation, due to the artificial binning.

In the **Supplemental Information**, we present an alternative binning method, referred to as the ‘credit system’ (outlined in **SI Figure S1 panel E & K, and Table S2 sub-heading 3**), where credits are allocated for extra keypresses at the end of a bin (for both groups), and at the beginning (only for the group with no breaks). This method offered an adjustment for the group with no breaks, whose sequences might have been interrupted by the artificial binning. The method yielded very similar results, as shown in **SI Table S2 sub-heading 3**.

In **Figure 1 panels C and E**, all 20-second test periods were divided into two 10-second bins, with the average number of correct keypresses across the two bins representing a single value for each 20-second period. This binning was also applied for statistical purposes, specifically to T1 and T2 when examining practice effects, where they serve as baselines for comparison with the binned training periods. This approach ensured consistency in keypress counting and accounted for any variations that could arise from the binning process. However, the primary statistical comparisons between groups at each of the five test periods are based on the total number of correct keypresses across the full 20-second test durations, without artificial binning.

#### Tapping speed of a sequence

Tapping speed of a completely correct sequence was evaluated as the inverse of the mean of the four inter-press intervals separating the five keypresses that constituted a fully correct sequence (keypresses per second).

#### Online and offline gains

Using the same method as Bönstrup et al.^1,2^, the difference in tapping speed between the first and the last correct sequence within a practice period was evaluated as ‘micro-online gains’, and the difference in tapping speed between the last correct sequence of a practice period and the first correct sequence of the next practice period was evaluated as ‘micro-offline gains’, i.e., MOG. The sum of these performance changes across all trials, results in the ‘online gain’ or ‘offline gain’ for each participant.

In Experiments 3, 4 and 5, we averaged the time between five consecutive responses that corresponded to a sequence (pre-defined by us as described above, but unknown to the participants), and obtained tapping speed for correct sequences by taking the inverse. The last five correct keypresses which were a part of a complete sequence were considered to be the last correct sequence of a practice period. For evaluation of the sum of offline performance improvements in both conditions of Experiment 5 (“Window size 5” or “Window size 1”), no MOG were computed for the rest periods for which the condition changed (i.e., every fourth rest period was omitted). In Experiments 3 and 4, the sum of performance improvements across 10 trials was used, whereas for Experiment 5, the sum across 15 trials were used, after discarding the transition trials.

#### Movement initiation time

This was evaluated as the time from the start of the practice period until the first response was made. Therefore, initiation times (**Figure 3E**) were evaluated only for the trials in which the first response was part of a fully correct sequence. The median across trials were then averaged (mean) across participants.

#### Statistical analysis

For statistical testing of online and offline performance changes, i.e. MOG, we computed one-sample t-tests against zero in all experiments. In Experiments 1 and 2, we compared learning between groups via 2×5 repeated measures ANOVA (two groups, five test sessions). For the two training blocks, we computed two 2×4 repeated measures ANOVA (two groups, four time points: baseline test and three training bins), one for each training block. The baseline for each training block was the average of performance in the two 10-second bins which were a part of the test session that preceded the respective training block (i.e., first test session for the first training block, and third test session for the second training block). The Greenhouse Geisser correction was used for all instances where the assumption of sphericity was violated. Interaction effects were followed up by t-tests. We corrected all p-values in these cases for multiple comparisons using the Holm-Bonferroni method. For statistical comparison of MOG between the ‘Non-repeating’ groups of Experiments 3 and 4, a Mann-Whitney U test was used to account for unequal variances between the groups of the two experiments, whereas an independent samples t-test was used for comparison of groups within each experiment. In Experiment 5, we used Pearson’s correlation to correlate inter-individual differences in tapping speed between conditions (“Window size 5” and “Window size 1”) with differences in movement initiation times. All statistical analyses were conducted in JASP (Version 0.17.1.0, jasp-stats.org) and MatLab (The Mathworks).

