## Supplemental Information for "“Micro-offline gains” convey no benefit for motor skill learning"

| bin | In-lab study | | | | | Online study | | | | |
|---|---|---|---|---|---|---|---|---|---|---|
| **1. Training sessions** | | | | | | | | | | |
| bin | *$BF_{10}$* | *t(83)* | *$p_{unc}$* | *$p_{corr}$* | *d* | *$BF_{10}$* | *t(356)* | *$p_{unc}$* | *$p_{corr}$* | *d* |
| T1 (mean 2 bins) | .281 | -0.705 | .483 | 1 | -0.153 | .703 | -1.935 | .054 | .216 | -0.205 |
| Training 1_bin1 | .342 | 0.971 | .335 | 1 | 0.211 | .438 | 1.660 | .098 | .294 | 0.175 |
| Training 1_bin2 | .609 | 1.508 | .135 | .540 | 0.327 | 129.38 | 3.849 | <.001 | .008 | 0.407 |
| Training 1_bin3 | 2.049 | 2.262 | .026 | .182 | 0.491 | 1109.2 | 4.411 | <.001 | .008 | 0.466 |
| T3 (mean 2 bins) | .229 | 0.168 | .867 | 1 | 0.037 | .163 | -0.831 | .407 | .62 | -0.088 |
| Training 2_bin1 | 2.848 | 2.429 | .017 | .136 | 0.527 | .192 | 1.017 | .31 | .62 | 0.107 |
| Training 2_bin2 | 1.413 | 2.059 | .043 | .258 | 0.447 | 1.845 | 2.403 | .017 | .085 | 0.254 |
| Training 2_bin3 | 1.01 | 1.858 | .067 | .335 | 0.403 | 6.828 | 2.922 | .004 | .024 | 0.309 |
| **2. Test sessions** | | | | | | | | | | |
| bin | *In-lab study* | | | | | *Online study* | | | | |
| | *$BF_{10}$* | *t(83)* | *$p_{unc}$* | *$p_{corr}$* | *d* | *$BF_{10}$* | *t(356)* | *$p_{unc}$* | *$p_{corr}$* | *d* |
| T1 (baseline) | .241 | -0.372 | .711 | 1 | 0.081 | .969 | -2.102 | .036 | .18 | -0.22 |
| T2 (end 1st tr.) | .34 | 0.965 | .338 | 1 | 0.209 | .174 | 0.91 | .364 | 1 | 0.096 |
| T3 (1st retention) | .227 | 0.07 | .944 | 1 | 0.015 | .148 | -0.706 | .481 | 1 | -0.08 |
| T4 (end 2nd tr.) | .248 | 0.455 | .650 | 1 | 0.099 | .161 | -0.813 | .417 | 1 | -0.09 |
| T5 (2nd retention) | .227 | 0.101 | .920 | 1 | 0.022 | .12 | -0.248 | .804 | .962 | -0.03 |

**Table S1. Experiments 1 (in-lab) & 2 (online), in reference to Figure 1 panels C and E: 1). Training sessions:** Results from independent-samples t-tests comparing the number of correct keypresses between groups during training (bins 1-3, separately for the first and second training block) and pre-training baseline (T1 and T3). A 2x4 ANOVA (two groups and four time points: baseline and three training bins) revealed a main effect of time point for the first training block (in-lab study: F(2.4,199.8)=78.545, p<.001, $\eta^2_{partial}$=0.486; online study: F(2.9,1036.9) = 189.8, p<.001, $\eta^2_{partial}$= 0.348), as well as the second training block (in-lab study: F(2.8,229.7) = 24.835, p<.001, $\eta^2_{partial}$= 0.23; online study: F(2.9,1029.6) = 58.703, p<.001, $\eta^2_{partial}$= 0.142). Additionally, there was a group by time point interaction, indicating that this improvement differed between groups, both for the first training block (in-lab study: F(2.4,199.8) = 7.604, p<.001, $\eta^2_{partial}$= 0.084; online study: F(2.9,1036.9) = 27.2, p<.001, $\eta^2_{partial}$=0.071), and the second training block (in-lab study: F(2.8,229.7) = 6.59, p<.001, $\eta^2_{partial}$= 0.074; online study: F(2.9,1029.6) = 17.482, p<.001, $\eta^2_{partial}$= 0.047). **2). Test sessions:** Results from independent-samples t-tests comparing the number of correct keypresses between groups in each of the five test sessions (T1 - T5). We corrected p-values for multiple comparisons using the Bonferroni-Holm method ($p_{corr}$; $p_{unc}$ represents uncorrected p-values). All tests reported in the table are two-sided.

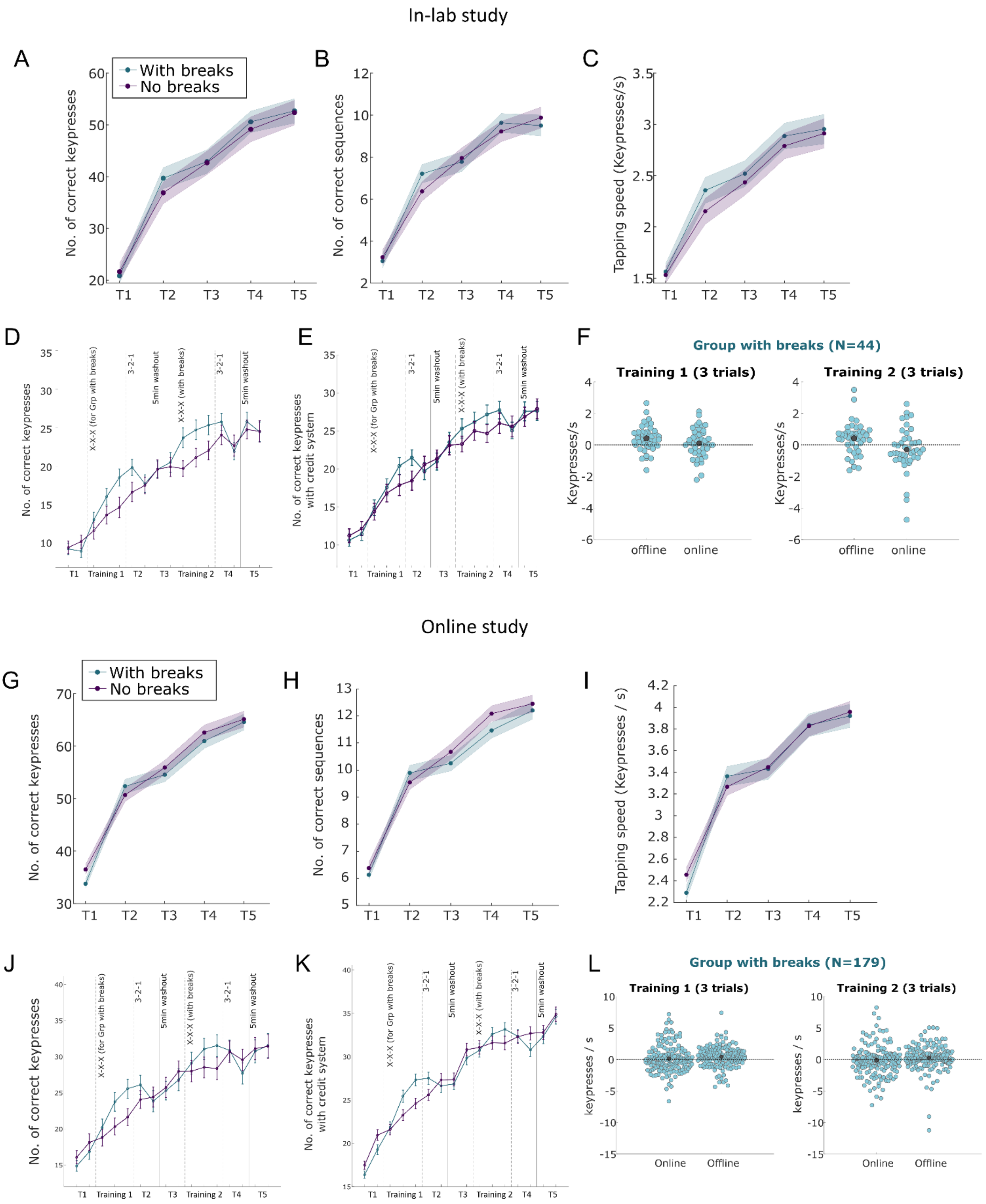


**Figure S1, panels A-F. In reference to data from Experiment 1 (In-lab study, With breaks group, n = 44 & No breaks group, n = 41).** Panels **A**, **B** and **C** display the number of correct keypresses, the number of correct sequences or the mean tapping speed of correct sequences, between the two groups across each 20-s test period. No significant differences were observed between the groups in any 20-s test period for these measures. Panels **D** and **E** present the number of correct keypresses for each 10-s bins (artificially binned) throughout the experiment, without (**D**) and with a credit system (**E**). Panel **F** shows the 'Micro-offline gains' for the group with breaks, during both training periods. **Figure S1, Panels G-L. In reference to data from Experiment 2 (Online study, With breaks group, n = 179 & No breaks group, n = 179).** Panels **G**, **H** and **I** display the number of correct keypresses, the number of correct sequences or the mean tapping speed of correct sequences, between the two groups across each 20-s test period. Panels **J** and **K** present the number of correct keypresses for each 10-s bins (artificially binned) throughout the experiment, without (**J**) and with a credit system (**K**). Panel **L** shows the 'Micro-offline gains' for the group with breaks, during both training periods.

1. Results of ANOVA and independent samples t-test for in-lab study (Figure S1, panels A, B and C)

| Effect | ***Number of correct keypresses*** | | | ***Number of correct sequences*** | | | ***Tapping speed*** | | |
|---|---|---|---|---|---|---|---|---|---|
| | *F* | *p* | $\eta^2_{partial}$ | *F* | *p* | $\eta^2_{partial}$ | *F* | *p* | $\eta^2_{partial}$ |
| Test | 250.1 | <.001 | .751 | 166.6 | <.001 | .667 | 170.492 | <.001 | 0.298 |
| Group | .093 | .761 | .001 | 0.041 | .841 | 4.9e-4 | .344 | .559 | 0.003 |
| Test*Group | .776 | .504 | .009 | 1.52 | .196 | .018 | .799 | .481 | 0.001 |

| Test # | ***Number of correct keypresses*** | | | ***Number of correct sequences*** | | | ***Tapping speed*** | | |
|---|---|---|---|---|---|---|---|---|---|
| | *t (83)* | $p_{unc}$ | *d* | *t (83)* | $p_{unc}$ | *d* | *t (83)* | $p_{unc}$ | *d* |
| T1 | -0.372 | .711 | -0.081 | -0.359 | .721 | -0.078 | 0.265 | .791 | 0.06 |
| T2 | 0.965 | .338 | 0.209 | 1.322 | .19 | 0.287 | 1.183 | .24 | 0.257 |
| T3 | 0.07 | .944 | 0.015 | -0.265 | .792 | -0.057 | 0.471 | .639 | 0.102 |
| T4 | 0.455 | .65 | 0.099 | 0.637 | .526 | 0.138 | 0.518 | .606 | 0.113 |
| T5 | 0.101 | .92 | 0.022 | -0.53 | .597 | -0.115 | 0.216 | .829 | 0.047 |

2. Results of ANOVA and independent samples t-test for in-lab study (Figure S1, panels G, H and I)

| Effect | ***Number of correct keypresses*** | | | ***Number of correct sequences*** | | | ***Tapping speed*** | | |
|---|---|---|---|---|---|---|---|---|---|
| | *F* | *p* | $\eta^2_{partial}$ | *F* | *p* | $\eta^2_{partial}$ | *F* | *p* | $\eta^2_{partial}$ |
| Test | 591.5 | <.001 | 0.624 | 466.5 | <.001 | 0.567 | 424.415 | <.001 | 0.546 |
| Group | .3 | .584 | 8.43e-4 | .482 | .488 | 0.001 | .029 | .864 | 8.27e-5 |
| Test*Group | 2.9 | .03 | 0.008 | 2.7 | .038 | 0.008 | 2.543 | .056 | 0.007 |

| Test # | ***Number of correct keypresses*** | | | ***Number of correct sequences*** | | | ***Tapping speed*** | | |
|---|---|---|---|---|---|---|---|---|---|
| | *t (356)* | $p_{unc}$ | *d* | *t (356)* | $p_{unc}$ | *d* | *T (365)* | $p_{unc}$ | *d* |
| T1 | -2.102 | .036 | -0.222 | -0.865 | .388 | -0.091 | -1.862 | .063 | -0.198 |
| T2 | 0.91 | .364 | 0.096 | 0.915 | .361 | 0.097 | 0.799 | .425 | 0.085 |
| T3 | -0.706 | .481 | -0.075 | -1.024 | .307 | -0.108 | -0.121 | .904 | -0.013 |
| T4 | -0.813 | .417 | -0.086 | -1.515 | .131 | -0.16 | 0.057 | .955 | 0.006 |
| T5 | -0.248 | .804 | -0.026 | -0.539 | .59 | -0.057 | -0.253 | .8 | -0.027 |

3. Results of independent samples t-test for in-lab and online studies, using credit system (Figure S1 panels E & K)

| **Time point** | ***In-lab study*** | | | ***Online study*** | | |
|---|---|---|---|---|---|---|
| | *t(83)* | $p_{unc}$ | *d* | *t(356)* | $p_{unc}$ | *d* |
| T1 (mean of 2 bins) | -0.579 | .564 | -0.126 | -2.089 | .037 | -0.221 |
| Training 1_bin1 | 0.314 | .755 | 0.068 | 0.285 | .776 | 0.03 |
| Training 2_bin2 | 0.429 | .669 | 0.093 | 2.354 | .019 | 0.249 |
| Training 3_bin3 | 1.444 | .153 | 0.313 | 2.967 | .003 | 0.314 |
| T3 (mean of 2 bins) | 0.012 | .99 | 0.003 | -0.744 | .457 | -0.079 |
| Training 2_bin1 | 1.242 | .218 | 0.27 | -0.313 | .755 | -0.033 |
| Training 2_bin2 | 0.641 | .523 | 0.139 | 0.927 | .354 | 0.098 |
| Training 2_bin3 | 1.461 | .148 | 0.317 | 1.458 | .146 | 0.154 |

4. Results for trainings separately, in-lab and online studies (Figure S1 panels F & L)

| Sum of improvements separately for trainings | ***In-lab study*** | | | | ***Online study*** | | | |
|---|---|---|---|---|---|---|---|---|
| | t (84) | $p_{unc}$ | d | *BF₁₀* | *t (357)* | $p_{unc}$ | *d* | $BF_{10}$ |
| Training 1 offline | 3.011 | .003 | 0.327 | 7.793 | 4.055 | <.001 | 0.214 | 172.125 |
| Training 2 offline | 1.875 | .064 | 0.203 | .636 | 1.322 | .187 | 0.07 | .141 |
| Training 1 online | 1.5 | .137 | 0.163 | .351 | 2.582 | .01 | 0.136 | 1.57 |
| Training 2 online | -0.525 | .601 | -0.057 | .137 | 0.645 | .519 | 0.034 | .073 |

**Table S2.** Sub-heading 1 (in reference to **Figure S1**, **panels A, B, and C**, Experiment 1) and Sub-heading 2 (in reference to **Figure S1, panels G, H, and I**, Experiment 2). For both sub-headings: Results of a 2 x 5 Repeated measures ANOVA for two groups and five 20-second test sessions, in the number of correct keypresses, the number of correct sequences and the mean tapping speed. **Below**: Results from independent-samples t-tests comparing the number of correct keypresses, the number of correct sequences and the mean tapping speed between the two groups at each of the five 20-second test sessions. **Sub-heading 3, in reference to Figure S1, panels E and K (Experiments 1 & 2).** Results from independent-samples t-tests comparing the number of correct keypresses between the two groups during training, using a credit system to account for interrupted sequences during the artificial binning (**Figure S1,** panel **E** and **K**). When the keypresses were artificially split into 10-second bins, sequences might be divided between bins. For example, a correctly typed 5-element sequence might be split, with 3 keypresses falling in one bin and 2 in the next. According to our method for evaluating correct keypresses (see Data Analysis in the main manuscript), keypresses at the boundary of a bin would not be counted if fewer than three consecutive keypresses matched the sequence. This could also occur at the start of a bin if the sequence did not begin with "4-1-3...". To address such cases, we implemented an additional credit system. In this system, any instance of 2 or fewer correctly ordered keypresses at the end of a bin (for both groups) was added to the total correct keypress count for that bin. Additionally, at the start of the next bin, for the 'No Breaks' group only, we counted keypresses that corresponded to correct sequences interrupted by artificial binning. We observed similar results regardless of the method used. A 2x4 ANOVA (two groups and four time points: baseline and three training bins) revealed a main effect of time point for the first training block (in-lab study: $F(2.6,215.9) = 77.353$, $p<.001$, $\eta^2_{partial} = 0.482$; online study: $F(3,1068) = 182.3$, $p<.001$, $\eta^2_{partial} = 0.339$), as well as the second training block (in-lab study: $F(3,249) = 20.419$, $p<.001$, $\eta^2_{partial} = 0.197$; online study: $F(2.9,1027.4) = 54.293$, $p<.001$, $\eta^2_{partial} = 0.132$). Additionally, there was a group by time point interaction, indicating that this improvement differed between groups, both for the first training block (in-lab study: $F(2.6,215.9) = 2.94$, $p=0.04$, $\eta^2_{partial} = 0.034$; online study: $F(3,1068) = 15.3$, $p<.001$, $\eta^2_{partial} = 0.041$), and the second training block for the online study ($F(2.9,1027.4) = 5.937$, $p<.001$, $\eta^2_{partial} = 0.016$), and a trend towards significance for the in-lab study ($F(3,249) = 2.182$, $p=0.091$, $\eta^2_{partial} = 0.026$). **Sub-heading 4, in reference to Figure S1, panels F and L (Experiments 1 & 2).** Results from one-sample t-tests against zero for the sum of online and offline performance improvements (i.e., micro online and offline gains) during each training separately, for the group with breaks (**Figure S1, panels F** and **L**). All tests reported in the table are two-sided ($p_{unc}$ represents uncorrected p-values).

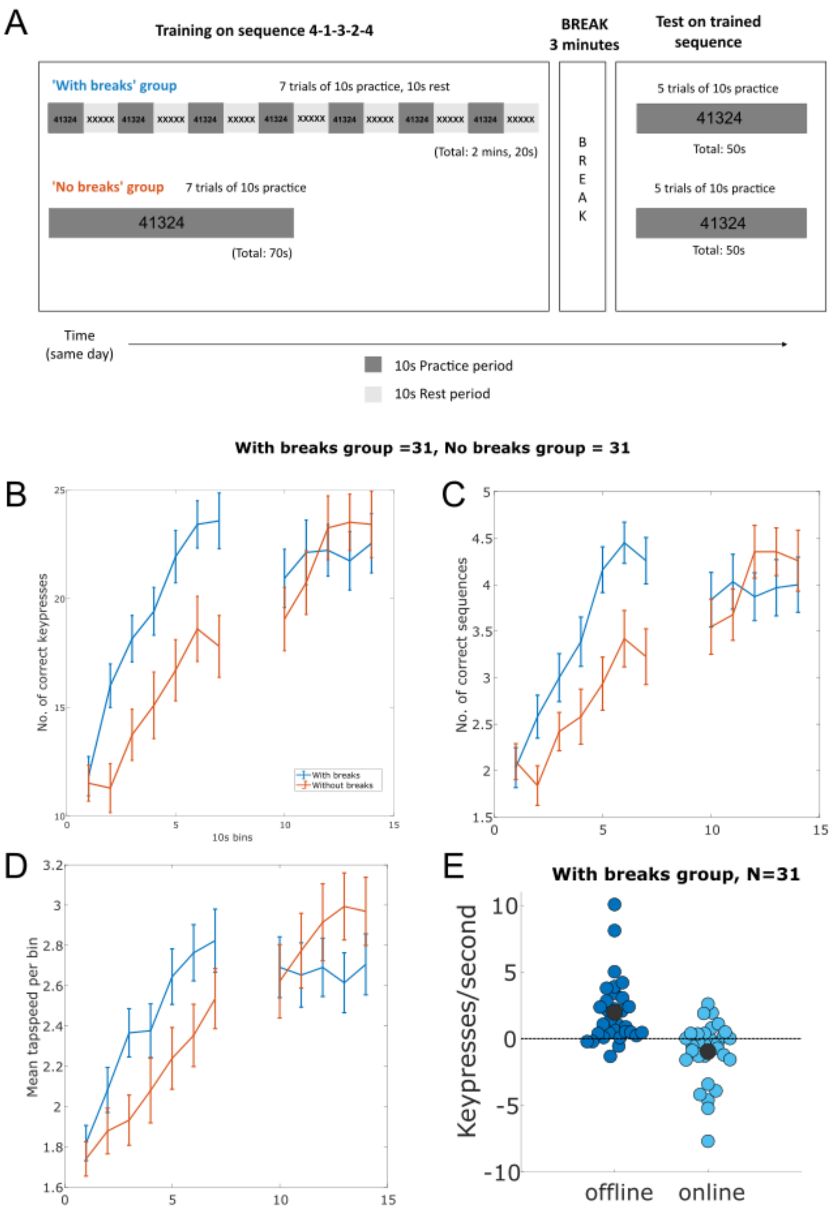


**Figure S2. Additional Experiment S1. Training with breaks vs no breaks.** Prior to running Experiments 1 and 2, we ran a similar study with n = 62 participants to test the impact of breaks during early motor skill acquisition (31 females, mean age 26.4 ± 3.5 years; With breaks, n = 31, No Breaks, n = 31). Panel **A** outlines the experiment design. In the group with breaks, participants practiced the sequence of finger movements 4-1-3-2-4 repeatedly through 7 trials, each lasting 10-s, followed by a 10-s rest. In contrast, the group without breaks practiced the sequence continuously for 70 seconds. Both groups then underwent a 3-minute washout period to eliminate fatigue or task-related effects. Skill retention was evaluated in a 50-s continuous practice block following the washout period. Participants were incentivized with bonus money based on the total number of correct sequences completed throughout the experiment. Analyses were performed in the same way as for the main experiments 1 & 2. We observed significant MOG in the group with breaks with superior performance in terms of correct keypresses during several 10-s intervals during training compared to the group without breaks. However, after the 3 minute washout, no group differences were observed during the 50-s test period. Panel **B** represents the number of correct keypresses, panel **C** represents the number of correct sequences and panel **D** presents the mean tapping speed of correct sequences, for both groups across training and test periods, in 10-s bins. **E** shows the micro-offline and micro-online gains during training for the group that trained with breaks. There were significant micro-offline gains (MOG) against zero ($t(30)=4.453$, $p<.001$, $d=0.8$, $BF_{10}=239.153$) but negative micro-online gains ($t(30)=-2.371$, $p<.024$, $d=-0.426$, $BF_{10}=2.118$) for the group that trained with breaks, across 7 trials (**Figure S2,** panel **E**). All error bars represent the SEM.

| 1. Training | | | | | | | | | |
|---|---|---|---|---|---|---|---|---|---|
| Effect | ***Number of correct keypresses*** | | | ***Number of correct sequences*** | | | ***Tapping speed per 10s*** | | |
| | *F (1,60)* | *p* | $\eta^2_{partial}$ | *F (1,60)* | *p* | $\eta^2_{partial}$ | *F (1,60)* | *p* | $\eta^2_{partial}$ |
| Training | 129.213 | <.001 | 0.683 | 86.574 | <.001 | .591 | 100.857 | <.001 | .66 |
| Group | 4.759 | .033 | 0.073 | 2.746 | .103 | .044 | 1.165 | .285 | .022 |
| Training*Group | 11.81 | .001 | 0.164 | 9.253 | .003 | .134 | 1.985 | .165 | .037 |

| 2. Test | | | | | | | | | |
|---|---|---|---|---|---|---|---|---|---|
| Effect | ***Number of correct keypresses*** | | | ***Number of correct sequences*** | | | ***Tapping speed per 10s*** | | |
| | *F (1,60)* | *p* | $\eta^2_{partial}$ | *F (1,60)* | *p* | $\eta^2_{partial}$ | *F (1,60)* | *p* | $\eta^2_{partial}$ |
| Test | 16.09 | <.001 | .211 | 6.877 | .011 | 0.103 | 6.384 | .014 | .099 |
| Group | .072 | .79 | .001 | .002 | .968 | 2.8e-5 | .209 | .65 | .004 |
| Test*Group | 3.397 | .07 | .054 | 2.726 | .104 | 0.043 | 4.58 | .037 | .073 |

| 3. Independent samples t-test comparisons | | | | | | | | | |
|---|---|---|---|---|---|---|---|---|---|
| Time point | ***Number of correct keypresses*** | | | ***Number of correct sequences*** | | | ***Tapping speed per 10s*** | | |
| | *t (60)* | $p_{unc}$ | *d* | *t (60)* | $p_{unc}$ | *d* | *t(60)* | $p_{unc}$ | *d* |
| Training bin 1 | 0.263 | .793 | 0.067 | -0.223 | .824 | -0.057 | 0.607 | .547 | 0.164 |
| Training bin 2 | 3.132 | .003 | 0.796 | 2.362 | .021 | 0.6 | 1.204 | .234 | 0.325 |
| Training bin 3 | 2.782 | .007 | 0.707 | 1.757 | .084 | 0.446 | 2.487 | .016 | 0.637 |
| Training bin 4 | 2.294 | .025 | 0.583 | 2.029 | .047 | 0.515 | 1.397 | .168 | 0.361 |
| Training bin 5 | 2.826 | .006 | 0.718 | 3.253 | .002 | 0.826 | 1.954 | .055 | 0.5 |
| Training bin 6 | 2.599 | .012 | 0.66 | 2.748 | .008 | 0.698 | 1.967 | .054 | 0.5 |
| Training bin 7 | 3.016 | .004 | 0.766 | 2.648 | .01 | 0.673 | 1.31 | .195 | 0.338 |
| Test bin 1 | 0.945 | .348 | 0.24 | 0.696 | .489 | 0.177 | 0.291 | .772 | 0.075 |
| Test bin 2 | 0.663 | .51 | 0.169 | 0.878 | .383 | 0.223 | -0.493 | .624 | -0.125 |
| Test bin 3 | -0.547 | .587 | -0.139 | -1.263 | .212 | -0.321 | -0.937 | .353 | -0.238 |
| Test bin 4 | -0.951 | .345 | -0.242 | -0.977 | .333 | -0.248 | -1.698 | .095 | -0.431 |
| Test bin 5 | -0.426 | .672 | -0.108 | -0.583 | .562 | -0.148 | -1.159 | .251 | -0.294 |

**Table S3**. **Experiment S1, in reference to test of Figure S2 panels B, C and D. Sub-heading 1.** Results of a 2 x 2 repeated measures ANOVA (2 groups and 2 trials: 1$^{st}$ and 7$^{th}$ 10-s bins), in the number of correct keypresses, the number of correct sequences and the mean tapping speed, for the training trials. **Sub-heading 2.** Results of a 2 x 2 repeated measures ANOVA (2 groups and 2 test bins: 1$^{st}$ and 5$^{th}$ 10-s bins), in the number of correct keypresses, the number of correct sequences and the mean tapping speed, for the test trials. **Sub-heading 3.** Results from independent-samples t-tests comparing the number of correct keypresses, the number of correct sequences and the mean tapping speed between the two groups for all 10-s bins (artificially binned throughout the experiment). All tests reported in the table are two-sided ($p_{unc}$ represents uncorrected p-values).

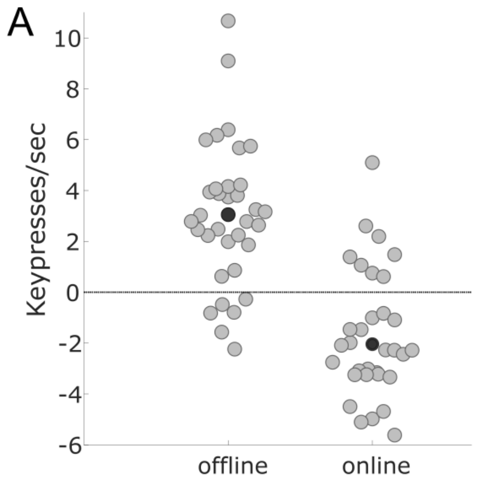


**Figure S3: Conceptual replication of Bönstrup et al.'s 2019 main paradigm**. Data from a conceptual replication using the same task paradigm (N=34, 16 females, mean age = 27.3 ± 4.3 years) as in *Bönstrup et al. 2019, Current Biology*[1]*[S1]*. There were 36 trials with 10-s practice periods interleaved by 10-s rest periods. The sequence used was 4-1-3-2-4 and the feedback for every keypress was presented in the same way as *Bönstrup et al. 2019.* Participants heard white noise throughout the experiment, in order to prevent distractions as well as any learning as a function of keypress sound from the keyboard. The results show significantly positive 'micro-offline gains' (t(33) = 6.307, p = <.001, d = 1.082, $BF_{10}$ = 41642.126) and significantly negative 'micro-online gains' (t(33) = -4.224, p = <.001, d = -0.724, $BF_{10}$ = 150.369), across the first 12 and 11 trials[1], respectively.
